# Computing and Optimizing Over All Fixed-Points of Discrete Systems on Large Networks

**DOI:** 10.1101/2020.03.04.960724

**Authors:** James R. Riehl, Maxwell I. Zimmerman, Matthew F. Singh, Gregory R. Bowman, ShiNung Ching

## Abstract

Equilibria, or fixed points, play an important role in dynamical systems across various domains, yet finding them can be computationally challenging. Here, we show how to efficiently compute all equilibrium points of discrete-valued, discrete-time systems on sparse networks. Using graph partitioning, we recursively decompose the original problem into a set of smaller, simpler problems that are easy to compute, and whose solutions combine to yield the full equilibrium set. This makes it possible to find the fixed points of systems on arbitrarily large networks meeting certain criteria. This approach can also be used without computing the full equilibrium set, which may grow very large in some cases. For example, one can use this method to check the existence and total number of equilibria, or to find equilibria that are optimal with respect to a given cost function. We demonstrate the potential capabilities of this approach with examples in two scientific domains: computing the number of fixed points in brain networks and finding the minimal energy conformations of lattice-based protein folding models.

## Introduction

A fundamental element in the study of physics and dynamical systems is the notion of fixed points or equilibria. In the absence of exogenous input or disturbance, a system will not deviate from a fixed point, and the dynamics in the neighborhood of that point determines its stability, i.e. whether it is an attractor, repeller, or other type of critical point. Indeed, the state of a system (for our purposes, defined on a network) evolves in time on a state space that is landmarked by equilibria. Knowledge of these equilibria can therefore help to predict long term evolution of system trajectories. It is often particularly useful to know the equilibria that are optimal with respect to some cost function, for example, to find ground states or minimum energy configurations of materials, protein sequences, or brain networks, or to determine the most cost efficient or beneficial states in social or economic networks.

We focus here on dynamical systems where the dependency structure of the components of the state is described by a graph, and each component can take one of a small number of values. These include simplified neural network models, models of decision or opinion dynamics on social and economic networks, evolutionary games on networks, and chemical reaction and protein folding models. The notion of an equilibrium or fixed point has wide-ranging interpretations across these applications. In neural network models, fixed points represent invariant patterns of activation that can be associated with memories, cognitive task goals, or default states of awareness [1]. In protein folding models, the fixed points correspond to physically realizable protein structures [2]. Equilibria in social networks can be interpreted as states in which all individuals are satisfied with their respective actions, opinions, or votes [3]. A tool for computing the fixed points in these systems would thus be of great consequence across scientific domains.

Unfortunately, even for discrete valued systems, finding all fixed points is computationally infeasible [4, 5]. All is not lost however, if the network under consideration has certain advantageous properties. In particular, if the network topology can be repeatedly partitioned by cutting no more than some fixed number (*k*) of edges, we say it is *k-separable*, and the approach we put forth in this paper can find the solution with a computation time that is linear in the size of the network and exponential only in local properties. When this holds for sufficiently small values of *k*, the proposed method can find all equilibria for very large systems, but even when it does not hold, we can solve the problem on much larger networks than with the more naive methods that are typically used in practice. Accordingly, the main contribution of this work is a provably tractable method for searching the global equilibrium space of discrete-valued systems on *k*-separable networks, using a recursive procedure of graph partitioning and sparse set operations, augmented by efficient bookkeeping. We emphasize that the algorithm yields the *exact* set of global fixed points in a sparse representation, from which individual equilibria can be queried according to some desired criteria, for example, those that minimize an energy function. We also show how to quickly check the existence and total number of equilibria, and explain how broader characterizations such as fixed-point landscapes can be obtained without generating the entire equilibrium set.

To highlight the practical versatility of this method, we provide two case studies. In the first, we analyze the dynamics of functional brain networks extracted from the Human Connectome Project database. We show that such networks exhibit exponential growth in the number of fixed points as a function of interconnected brain regions, a prediction made in theoretical studies but here verified empirically for the first time. In the second case study, we address the decades-old challenge of finding low-energy configurations of proteins, toward facilitating a deeper understanding of protein evolution and design. We consider a prototypical protein lattice model and are able to efficiently find the global energy minima as well as perform an exact characterization of the ruggedness of the fixed-point landscape. Each of these case studies features a stochastic real world system for which a deterministic system is constructed and used as an analytical tool. Specifically, we define the deterministic system such that its fixed points contain the set of energy extrema of the corresponding stochastic system.

## Discrete systems and their fixed points

We consider discrete-time and discrete-valued dynamical systems in the general form

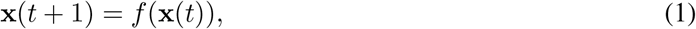

with state vector **x**(*t*):= [*x*_1_(*t*), …, *x*_*n*_(*t*)]^T^. Each component of the state *x*_*i*_(*t*) can take one of a finite set 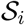 of possible values. Define a directed graph *𝔾* = (*𝒱*_0_, *ɛ*_0_) in which each node in *𝒱*_0_ corresponds to a component of the state, and the edges *ɛ*_0_ define dependencies in the system dynamics. Namely, an edge (*i, j*) ∈ *ɛ*_0_ indicates that *x*_*i*_(*t* + 1) may depend on *x*_*j*_(*t*). Conversely, the lack of an edge means there is no dependency. The dynamics are therefore separable into the components *x*_*i*_(*t* + 1) = *f*_*i*_(**x**(*t*)), in which *f*_*i*_ only depends on the states of node *i* and its neighbors in the network.

A fixed point or *equilibrium* **x**^*\**^of the system is a state in which no node will change between time steps, i.e. *f* (**x**^*\**^) = **x**^*\**^. Let Ω(*𝒱*_0_) denote the set of all such equilibria. Since *f* (**x**) is in general nonlinear, computing Ω(*𝒱*_0_) is a challenging task, and the main contribution of this work is a method for exploiting the network topology and local dynamics to compute a sparse representation of Ω(*𝒱*_0_) in an efficient way.

## Results

Our main result is an algorithm that computes and optimizes over the global equilibrium set of system (1). In this section, we describe how to do this using a combination of local dynamical analysis and recursive graph partitioning, discuss the computational complexity and related literature, and demonstrate the approach on practical applications in biochemistry and computational neuroscience.

### Finding and optimizing over all fixed points

The approach begins by computing the set of *local equilibrium* (LEQ) states for each node in the network. For a given node *i*, we define the LEQ as the set of all system states **x** such that *x*_*i*_(*t*) will not change at time *t* + 1 according to the local update rule *f*_*i*_(**x**(*t*)):

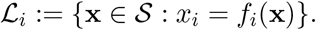

For sparsely connected networks in which the numbers of state values 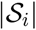 is not too large, these sets are fast to compute and can be efficiently stored using the sparse representations described in **Methods**. An important observation is that the global equilibrium set is equal to the intersection of all LEQ sets. Unfortunately, computing all of these intersections may be too difficult for large networks. However, by using graph partitioning while retaining necessary information for each partitioned region, we can decompose the problem in a way that allows for exact construction of the full set of equilibrium points (see **Methods: Recursive partition-based intersections**). Specifically, by intersecting the LEQ sets within each partitioned subnetwork, we form a set of *regional equilibrium* (REQ) states. This effectively reduces the problem to finding all compatible REQ states on the partitioned subnetworks, which can be achieved by intersecting neighboring LEQ sets if the subnetworks are small enough, or otherwise by performing an additional partition and proceeding recursively. The end result is a top-level REQ set along with a hierarchical data structure 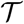 that efficiently encodes all information necessary to generate the full equilibrium set Ω(*𝒱*_0_).

What renders this algorithm tractable for *k*-separable networks is both the recursive decomposition and the fact that we need only a compact representation of the sets (including only the nodes on the partition boundaries) in order to construct higher level REQ sets and ultimately the global equilibrium set. The computational complexity thus depends primarily on the node degrees (number of connections to each node) and the number of edges that cross partition boundaries. Another key element in the approach is the construction of maps between the various stages of partitioning, which allow for quick access to the hierarchy of global, regional, and local equilibrium sets. Finally, if an energy or cost function is provided, the algorithm computes costs at each stage, which can be used to efficiently extract the optimal equilibrium states with respect to the given energy function. See **Methods** for a detailed presentation of this approach.

### Computational complexity

Special cases of finding all fixed points of system (1) such as deciding on the existence of a Nash equilibrium for certain systems [5] and calculating the minimum-energy state of ferromagnetic spin models [4] are proven to be *NP-hard*. This generally means that the time required by any existing algorithm to compute a solution is exponential in the size of the problem (e.g., number of nodes in the network). For example, the naive approach of checking for fixed points among all possible configurations in a network in which nodes can take one of two states would require a time proportional to 2^*n*^, and it is often not trivial to improve much on this brute force technique. When approximation is sufficient, methods such as simulated annealing [6, 7] and genetic algorithms [8, 9] can be effective in cleverly exploring the enormous state space, but these algorithms can generally only provide probabilistic guarantees on finding fixed points or global extrema. Otherwise, branch-and-bound algorithms [10] can reduce computation time by safely eliminating large regions of the total search space without compromising optimality. However, these methods often require significant effort in tailoring to a specific problem and their objective is typically to find a single global extremum. A recent approach to finding the global energy minimum on Ising lattices involves pairing combinatorial with convex optimization to converge upper and lower bounds on the solution, and was demonstrated on three-dimensional lattices containing up to 50 nodes, each of which can take two states [11]. In comparison, the method proposed here found the global minima and all fixed points on 3-dimensional lattices of 64 nodes, each of which can take three states (see **Case Study 2)**, and is demonstrated in on other types of networks having hundreds and thousands of nodes (see Fig. 1).

**Figure 1:**
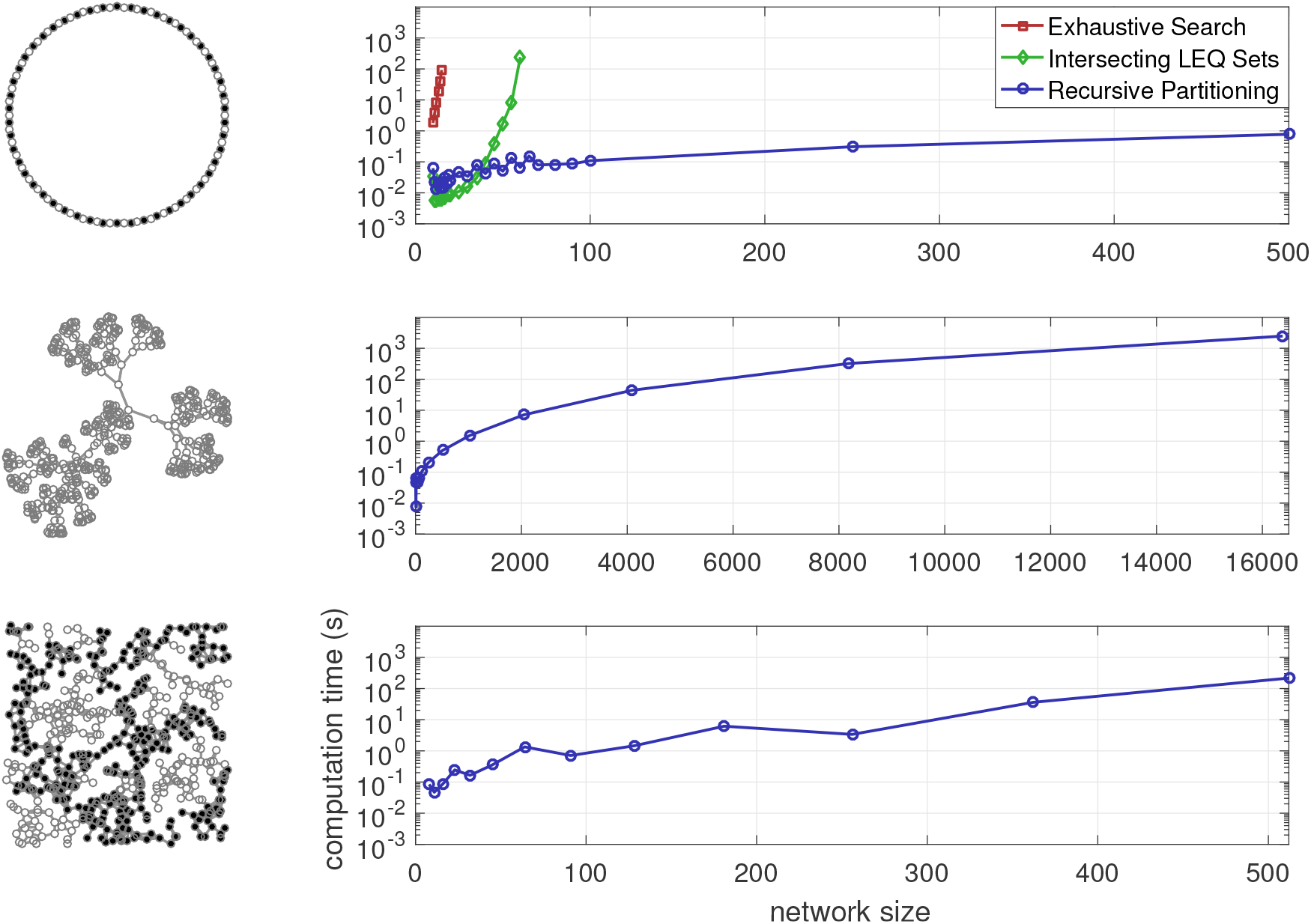
Time required to compute energy-extremal fixed points for three different network types: ring (top), tree (middle), and geometric random network (bottom). Node states are governed by linear threshold dynamics and energy is evaluated using a generalized Ising model (see **Supplementary Information (SI): Best-response and linear threshold dynamics**). Computation times for exhaustive search and intersecting all LEQ sets on ring networks are shown in the top panel. Black and white node colorings in the network diagrams respectively depict the states −1 and +1 in energy-extremal fixed points.

Significant effort has also been put toward finding attractors in *Boolean networks*, in which nodes take one of two values (0 or 1) and local dynamics are based on logic rules. This work is most closely associated with genetic regulatory network models, where nodes represent genes that are either expressed (1) or not (0), and edges indicate causal links between genes. Indeed, for the special case when each set 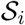 contains exactly two elements, system (1) is equivalent to a Boolean network. Several approaches have been proposed to find attractors in Boolean networks, including simulation [12], aggregation [13], binary decision diagrams [14], and satisfiability algorithms [15]. In particular, satisfiability rules paired with a partition of the Boolean network are used in [16] to find attractors in sequential and parallel implementations. We emphasize here three conceptual advancements over previous aggregation or partition-based techniques for finding attractors [13, 16]: (i) one level of partitioning may not be sufficient to make computation on large networks tractable – the extension to arbitrary levels of recursive partitioning yields what is to our knowledge the first provably tractable algorithm for computing fixed points on a particular class of networks, namely, those with local connectivity structure captured by the measure we call partition separability; (ii) computing the energy function at each stage of the decomposition allows for useful characterizations of the attractor landscape without explicitly computing the entire set of attractors, which may become prohibitively large; (iii) extending to more general discrete-valued networks makes it possible to analyze networks in which node states can take more than two values.

The key characteristic in determining how much advantage will be gained by using this approach is how easily the network can be decomposed into smaller parts. We define the *partition separability* of a network as the maximum number of edges that cross partition boundaries over all partitions used in the decomposition. For network topologies such that the partition separability is independent of network size, the computation time of the proposed algorithm is exponential only in local network properties, namely, the number of local connections or *degree* of the nodes. The top panel in Fig. 1 shows the substantial reduction in computational complexity when using the recursive partitioning approach compared to exhaustive search or intersecting all LEQ sets to compute all fixed points in networks governed by two-state best-response dynamics (See **SI: Best-response and linear threshold dynamics**). We note that the local intersection method is already a steep improvement over exhaustive search. While ring networks are particularly well-suited to this approach since they are 2-separable, perhaps even better suited are tree networks, which are 1-separable. Computation times for optimizing over the entire equilibrium space on tree networks are shown in the middle panel of Fig. 1.

Indeed, we can prove that the algorithm is tractable for *k*-separable networks. Let 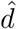 denote the maximum degree and *ŝ* the maximum number of local states 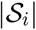 for any node *i* ∈ *𝒱*_0_ in the network, while *q* denotes the number of partition groups at each level, and *r* is the maximum number of nodes in a group at the lowest level of partitioning. The following theorem provides the parameterized computational complexity of the proposed algorithm, namely, that it is linear in the size of the network and exponential only in the node degree and the partition separability of the network (proof in **Appendix**).

#### Theorem 1.

*The complexity of computing the top-level REQ set* Ω^*p*^(*𝒱*_0_) *and the results tree* 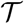 *for a k-separable network is at most O*(*αn*), *where* 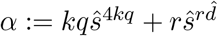.

For other types of networks, the algorithm still faces the inevitable hardness constraint, but the complexity is still reduced to the degree and separability properties, allowing solutions on much larger networks than possible through more naive methods. For example, we can find all fixed points of these dynamics on geometric random networks, such as in the bottom panel of Fig. 1, provided they are sufficiently sparse and separable. However, some other standard graph topologies are less well-suited to this approach. For example, since Erdos-Renyi random graphs have no local connectivity bias, the partition separability will always grow with the size of the network, and while scale free networks might have some local connectivity, the large degrees of hub nodes can be prohibitive even for the calculation of local equilibrium states. Determining the partition separability of a given network is itself a computationally complex problem, but there exist fast graph partitioning algorithms, e.g. [17, 18], which can be used to approximate this quantity. See **SI: Computational complexity analysis** for a comparison of the separability of several different network types, as well as a detailed analysis and proof of the computational complexity.

## Case Study 1: Finding the critical points in brain networks

A long-standing model of neural networks endows each neuron with a binary state representing whether it is active (firing) or inactive (not firing) [19]. Edges in such a model represent synaptic connections, which can be either excitatory, i.e. the firing of one neuron causes another to be more likely to fire, or inhibitory, i.e. the firing of one neuron causes another to be less likely to fire. Activation of a neuron is determined by a threshold on its presynaptic (incoming) activity.

In theoretical neuroscience, this class of model at the neuronal scale was initially used as a simplified descriptor of associative memories [1], where each fixed point in the system represents a complex but stationary pattern of neural activity corresponding to a particular memory, and, in the case of attractors, the region of attraction around the fixed point determines how easily the memory is recalled. Later work has examined stochastic variants of this model, closely related to the Ising model, to assess the role of correlations within neural populations and their effect on neural coding and information processing [20]. More recently, such models have also been used as the basis of theoretical studies to show how properties of neuronal networks might arise as a consequence of associative learning, wherein associations are encoded as equilibria of the underlying networks (27). The notion of energy via the Ising Hamiltonian is central to these analyses.

However, despite the relative simplicity of this model, many contemporary approaches rely on simulation studies and post-hoc statistical analyses [21]. Because the number of parameters in the model increases quickly with network size, such simulation-based approaches are inherently limited. Effort is thus being directed towards methods that can simplify the analyses of such models, especially as it pertains the model dynamics [22].

The overwhelming majority of functional network characterizations rely on static, graphical descriptions based on calculation of the pairwise correlation of the activity over a set of brain regions. Using dynamical systems models at such scales offers the potential for greater explanatory power relative to purely statistical descriptions [23, 24]. Such models span different spatiotemporal levels of description, from detailed biophysical models at neuronal scales, to mesoscopic, mean-field approximations at the level of brain regions. Often, these models incur a tradeoff between model complexity (including physiological interpretability/abstraction) and analytical tractability. The Ising framework provides a more abstract description of brain network activation, and has primarily been used as a means of analysis, especially in regards to establishing correlation between brain regions. However, the model itself does carry dynamical interpretability and recent efforts have highlighted its potential to generate hypotheses regarding the dynamical underpinnings of observed statistical outputs. For example, [25] uses the Ising framework to suggest how the architecture of a whole-brain scale network might impact its ‘repertoire’ of achievable states, as encoded through the network’s equilibria. Such model-based analysis and hypothesis generation is in the spirit of what we highlight in our Case Study. Indeed, an emerging area of study in functional neuroimaging pertains to nonstationary fluctuations in network covariance [26]. At a dynamical level, such fluctuations are fundamentally mediated by the attractor landscape of the system. However, given the high dimensionality, finding and characterizing all attractors is generally intractable. Here, as a first step to an attractor landscape characterization, we quantify the fixed points of 783 functional brain networks from the Human Connectome Project by ascribing linear threshold dynamics to the functional connectivity networks (see **Methods: Processing of data from Human Connectome Project**).

We observe in Fig. 2 that the number of fixed points increases exponentially with network size, with an exponent that varies for different resting state networks (subnetworks attributed to particular task specializations). In fact, the largest exponent was found in the frontoparietal network, suggestive of a large encoding capacity relative to networks such as a the ventral attention network, which possesses fewer stationary states. The frontoparietal network in particular is thought to be a key mediator of cognitive control [27, 28], though the extent to which its dynamics facilitate such is as yet opaque. Though our case study is an exploratory proof of concept, these results are intriguing as they suggest that this network may have a favorable dynamical landscape for encoding a diversity of cognitive states.

**Figure 2:**
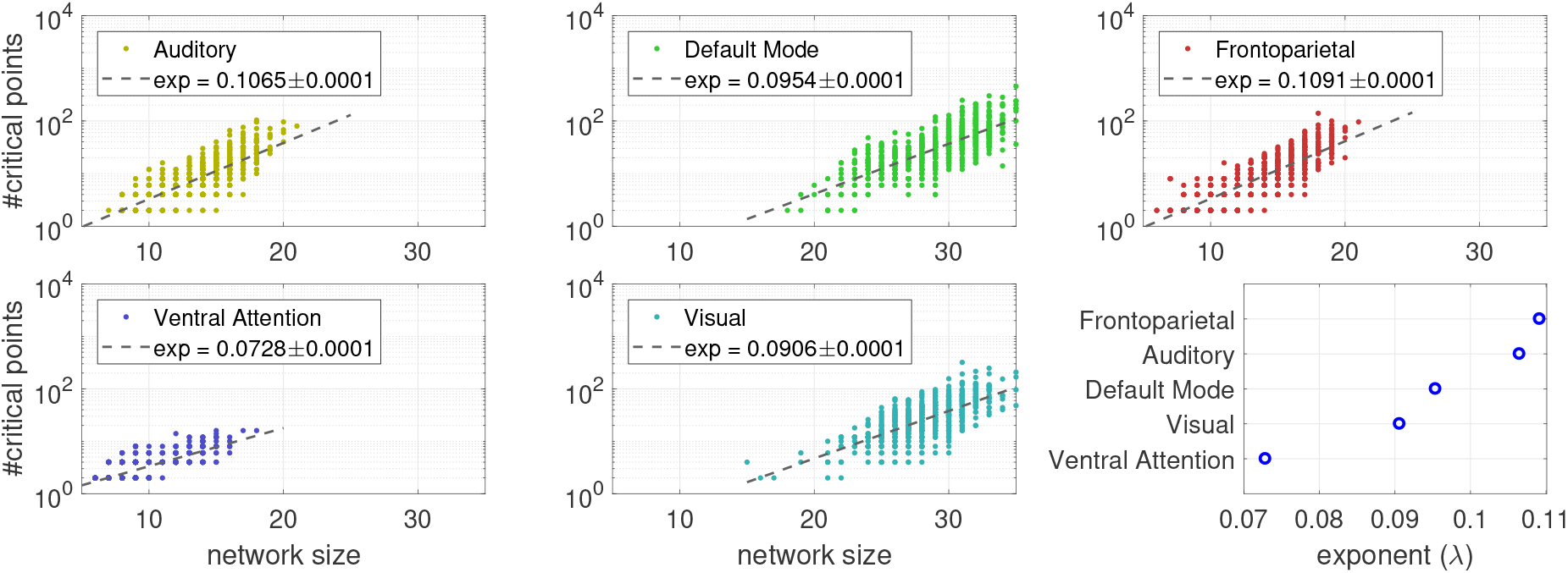
The number of critical points of discrete dynamical systems constructed from resting-state functional brain networks grows exponentially with network size, with an exponent that varies over the different resting state networks. The data comes from 783 participants in the Human Connectome Project (see **Methods**).

## Case Study 2: Analysis of a coarse-grained lattice model for protein folding

The biological function of a protein is highly dependent on its structure, i.e. how the linear sequence of amino acids folds onto itself in 3D space. While this can occur in many different ways, it is well-established that the most probable structures are those that minimize free energy, which is a function of hydrophobic and other intramolecular forces. Although a real protein sequence is composed of 20 amino acids, a reduced-order model can provide valuable insight for protein analysis and design. In particular, we use a model with 3 types of amino acids or residues: those that have positive charge, those that have negative charge, and those in which hydrophobic forces dominate. We consider a 3D lattice structure in which the nodes take values 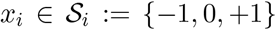, subject to the following energy function, which is a modification of the Hydrophobic Polar model, first introduced in [29]:

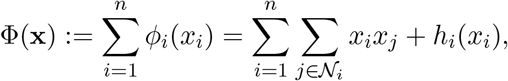

where 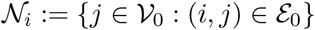 for each *i* ∈ *𝒱*_0_ and

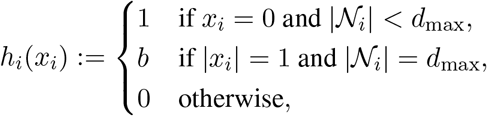

where *d*_max_ denotes the maximum node degree and *b* is a model parameter. We define the dynamics of the system such that at each time *k*, a random node updates to the state that minimizes the local energy *ϕ*_*i*_(*x*_*i*_). The point of this is not to directly model any physical dynamics, which are inherently stochastic, but rather to construct a deterministic system in which the set of fixed points is equivalent to the set of local minima of the energy function. We used the proposed algorithm to find all fixed points of this model for two different cases: a cubic lattice of dimension 4 *×* 4 *×* 4, and a more irregular structure also having 64 nodes, with *b* = 10. Since each node can take three values, there are a total of 3^64^ possible configurations in both models. For the lattice model, we find that 587,636,902 are fixed points (a fraction 1.7 *×* 10^*−*22^ of the total), and of these, only two are global minima. For the irregular model, there are 60,594,240 fixed points (a fraction of 1.8 *×* 10^*−*23^), 32 of which are global minima. Both optimal folding sequences for the regular lattice are depicted in Fig. 3, and the 32 global minima for the irregular model are shown in terms of eight independent components in Fig. 4. We can also plot the energy distributions of all fixed points for both models, as shown in Fig. 5. Perhaps counterintuitively, we observe a broader and more rugged fixed-point landscape in the cubic lattice than in the irregular lattice.

Note that cubic lattice structures are not *k*-separable for some constant value of *k* as the dimensions grow; rather, they are *s*^2^-separable, where *s* denotes the dimension of one side. As a result, regular lattice structures are not ideally suited to this approach. The advantage of this method becomes more striking for structures with some irregularities, in which the network topology is more suitable for partitioning. Indeed, the energy distribution of the irregular lattice was computed in approximately 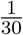 the time needed for the cubic lattice.

**Figure 3:**
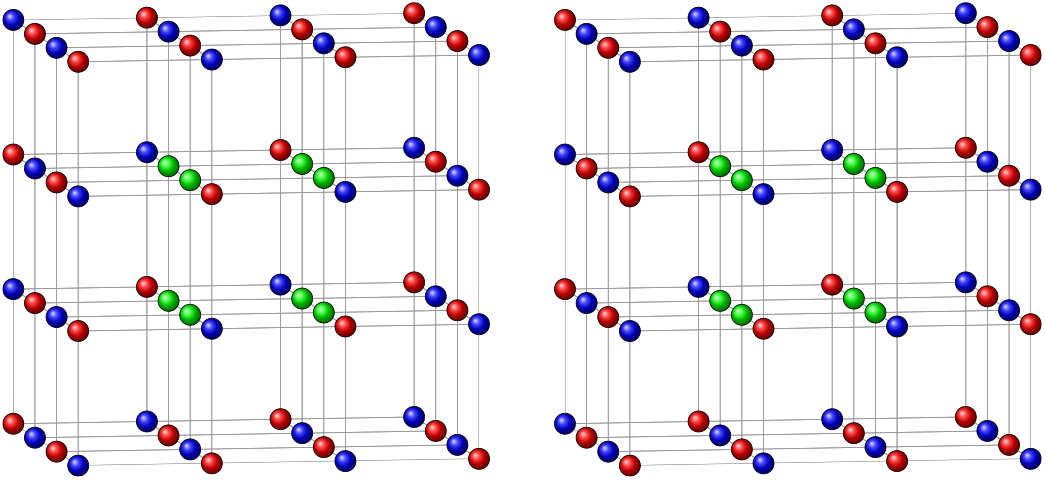
Optimal protein configurations on a 4 × 4 × 4 lattice. Blue and red indicate positive and negative charges, respectively, and green corresponds to hydrophobic residues.

**Figure 4:**
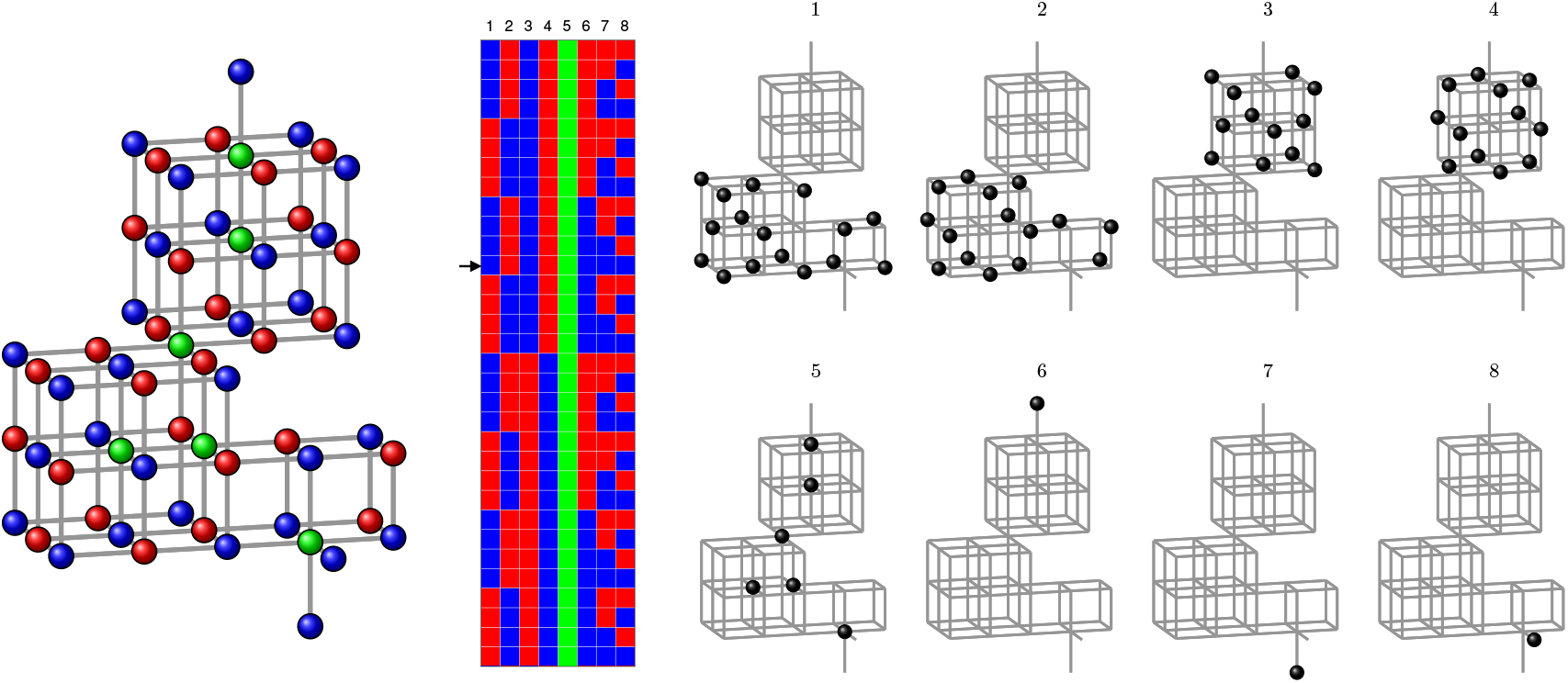
The 32 global energy minima in the irregular model, represented in the 32 rows of the color matrix, can be decomposed into 8 components, the nodes of which always take the same residue type, indicated by the color of the corresponding column entry. For example, the configuration shown on the left appears in row 12 of the color matrix, indicated by the arrow.

**Figure 5:**
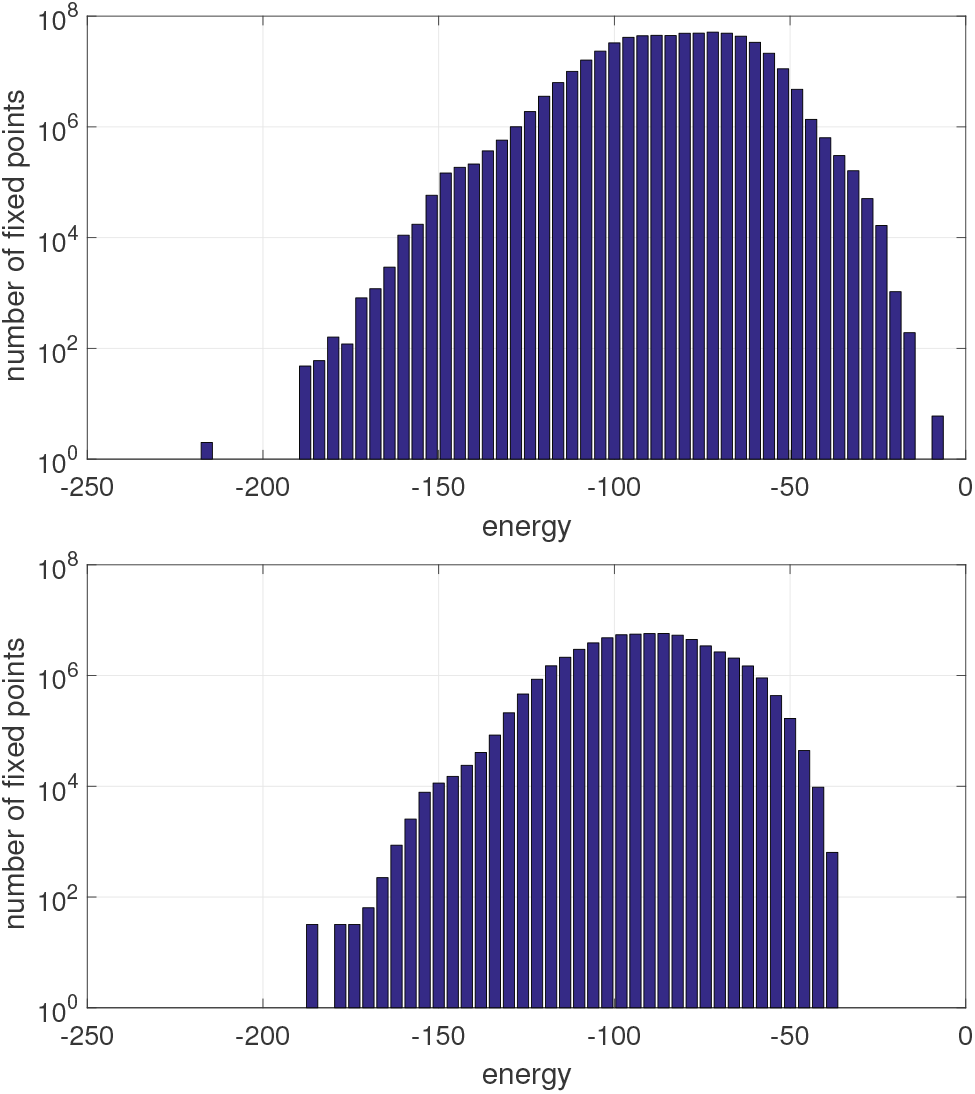
The distribution of energy minima in the cubic lattice (top) is both broader and more rugged than that of the irregular lattice (bottom).

Despite its simple appearance, the lattice model captures an essential challenge in protein folding and design: tackling sequence space. Even on this simplified model, the complexity of the problem is sufficient that an exhaustive search through sequence space is not possible. The standard practice for accessing sequence space relies on Monte Carlo searches, which fail to capture a vast majority of the fixed points for a given topology [30]. In the case of protein design, vast computational resources are spent sampling sequence space with the hopes of finding a sufficiently low local energy minimum. The proposed approach has the potential to significantly enhance this effort by providing a tractable means to map entire fixed-point landscapes, to within limits on protein size (see also **Discussion**).

### In the discussion of Case Study 2

As previously remarked, finding the fixed points of a system does not directly provide any information about convergence or stability. However, when the dynamics are known to produce trajectories that are decreasing with respect to the energy function in the neighborhood of particular fixed points, then Lyapunov theory predicts that these fixed points will indeed be stable attractors.

## Discussion

### Scientific Impact

The ability to find and optimize over all fixed points in discrete systems on networks has the potential to assist and advance research across several scientific domains.

Finding critical points of generalized Ising models is a long-standing problem in theoretical physics [31] that has remained active over the years [32, 11]. Expanding the scale and types of structures for which we can completely characterize the critical points has value in the study of magnetism, phase transitions, fluid dynamics, and other topics [33].

In biochemistry, finding the minimum energy protein sequence is a sought after task for which extraordinary computational resources and schemes have been devoted [34, 35]. Moreover, the ability to find all critical points can contribute to a greater understanding of the protein folding process, another target of large-scale computational projects [36]. Comparing the structure of proteins to the underlying fixed-point landscape can also give clues about the evolution of a protein sequence and provide greater insight into the impact of protein topology on evolutionary pathways.

Certain analyses in computational and theoretical neuroscience can be enabled through characterization of equilibria of (brain) network models. This is perhaps most prevalent in the study of memory encoding, where attractor networks have long been a popular model for the formation and recall of memoranda and other types of associations [37, 38]. More recently, notions of free energy minimization have been used as a theoretical schema within which to understand brain function at multiple spatial scales [39]. Testing such hypotheses has been difficult, due to the taxing nature of estimating energy landscapes from data. Perhaps as a result, the state of the art in brain network analyses, especially at whole-brain scales, has been largely limited to analyses of inter-region correlation and subsequent (static) graph theoretical analyses [40]. Nonetheless, key efforts in computational modeling and analysis are now underway to bridge architectural characterizations of brain networks to more faithful understanding of temporal dynamics over such networks [41, 42]. As shown here, the proposed methods can enable novel assessments of how brain network dynamics – via attractor and energy landscapes, which ultimately mediate the input to output lability of these networks [43] – are altered during cognition.

### Limitations

The methods presented in this paper provide significant advances in the ability to computationally characterize the dynamics of complex networked systems with discrete state spaces. In the context of whole-brain modeling (Case Study 2), the discrete-state model may be informative on second-to-second time-scales, upon which populations can be thought of as being active or not. Here, however, the coarseness of the state space precludes analysis over finer time-scales, when brain regions undergo more nuanced patterns of activation. Findings such as those we presented in Case Study 2 should thus be interpreted at the appropriate spatial and temporal scale, and not as a characterization of dynamics over continuous state spaces. Because our proposed approach is not restricted to binary models, somewhat richer analyses could be performed by augmenting the Ising-framework with additional (discrete) states.

Thus, from a practical standpoint, the proposed method will perform best for systems that can be modeled with a few discrete states per unit and with relatively sparse connection structure. When either the number of discrete states or node degrees become large, as in the case of scale-free networks for example, then even the first step of computing the local equilibrium sets becomes intractable. For example, analysis of a full 20-amino-acid protein-folding model or a micro-scale model of brain activity, where neurons have thousands of synaptic connections, would currently lie outside the capabilities of the proposed algorithm as implemented on standard personal computers.

## Methods

### Sparse set notations and operations

It is convenient to represent a subset of the state space 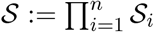 as the set of global states in which some of the nodes take specified values while others are allowed to take any value. This can be expressed as a sparse vector or a set of (*index, value*) pairs. For example, define **a** = {**x** ∈ {−1, 1}^5^: *x*_3_ = 1, *x*_5_ = *−*1}. We could alternatively denote this as 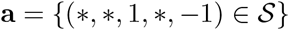 in which *\** indicates that an entry may take any value. Although illustrative, this notation may become cumbersome for higher dimensional systems, so we introduce the following more compact sparse representation:

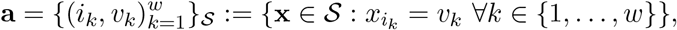

where *w* is the number of nodes whose state is defined. According to this notation, the example above would be expressed as 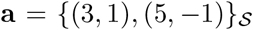, where 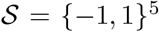. We also use the notation **a**[*i*] to access the defined element *i* ∈ *𝒱*_**a**_, where *𝒱*_**a**_ denotes the set of all nodes assigned in **a**. In the example, **a**[3] = 1 and **a**[5] = *−*1. We will achieve our objective primarily through the use of intersections and unions of these sparse representations of large subsets of the state space 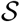. The following example demonstrates these operations on two sparsely represented subsets **a**, **b** ⊂ {−1, 1}^5^.

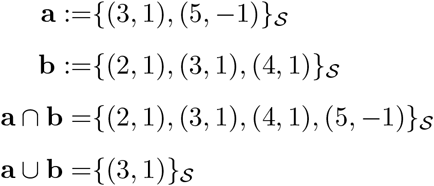

We see here that computing the union of two sets is achieved by performing an intersection of the sparse representations and vice versa, which allows for efficient computation of these operations.

We sometimes need to represent ordered sets of these sparsely represented subsets. For example, define 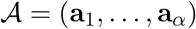 and 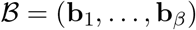 for some arbitrary integers *α* and *β*. Commonly used operations on these sets include the set of all pairwise intersections and the set of all pairwise unions, which we respectively denote by

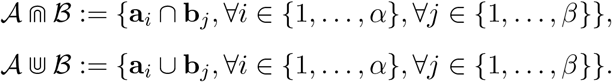

To denote the set of all *m*-way intersections and unions, we write

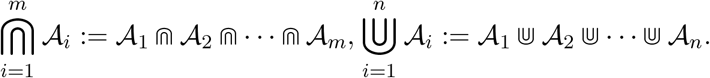

It is straightforward to confirm that the commutative, associative, and distributive properties hold for these operations. It will also be convenient to relate whether or not elements of an ordered set 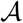 are subsets of elements of 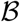. For this purpose, we define the matrix of size *α × β*:

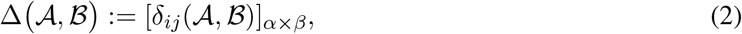

in which each entry is given by

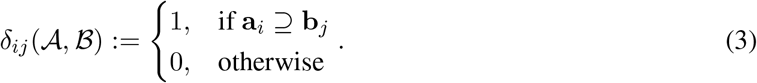

In addition, let 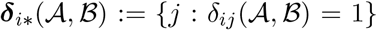 denote the index set of elements of 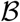 that are contained in **a**_*i*_, and let 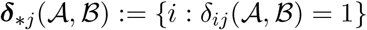 denote the index set of elements of 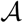 that contain **b**_*j*_.

### Recursive partition-based intersections

The next step after computing the LEQ sets (as described in **Results: Finding and optimizing over all fixed points**) is to partition the network into several smaller subnetworks and compute the regional equilibria on those subnetworks. The corresponding REQ sets can then be intersected using only the states of their boundary nodes to obtain a sparse representation of the global equilibrium set. At the end, we have all the information needed to quickly access any equilibrium without explicitly computing the whole set.

This method relies heavily on graph partitioning, which is itself a complex problem and an active research area. Although any valid graph partitioning algorithm can be used here, the choice may have a significant effect on performance. For example, an algorithm that tries to balance subgroup sizes while minimizing the number of edges that cross partition boundaries will generally lead to faster computation times than one that does not. It is also computationally advantageous for our purposes if the nodes on the partition boundaries have small LEQ sets. Optimally balanced graph partitioning is itself a computationally difficult problem. Fortunately, we do not require an optimal partition; rather, a reasonably good approximation will suffice, and recent research has shown the existence of algorithms that are tractable with respect to a parameterization of the requirements[44]. Indeed there are several approaches to choose from that are fast to compute relative to finding the fixed points. In our case studies and simulations, we used k-means clustering on the eigenvectors of the weighted adjacency matrix, with weights proportional to the size of the LEQ sets, which is a type of spectral graph partitioning algorithm [17].

Before proceeding, we introduce some definitions and notations related to partitioning the network. Let 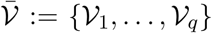 denote a *partition* of the node set *𝒱 ⊆ 𝒱*_0_ into *q* subsets, such that 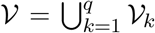 and *𝒱*_*j*_ ∩ *𝒱*_*k*_ = *∅* for all *j, k* ∈ {1, …, *q*}, *j ≠ k*. Further, we define the set of external boundary nodes 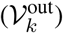 as the set of nodes that are not in *𝒱*_*k*_ but are adjacent (either to or from) a node in *𝒱*_*k*_. In contrast, we define the internal boundary nodes 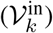 as the set of nodes that are in *𝒱*_*k*_ and are adjacent to a node that is not in *𝒱*_*k*_. We also define the union of these two sets as the set of combined boundary nodes 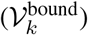. Formally, these sets are defined as follows:

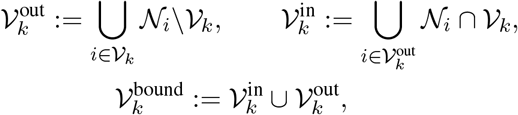

where 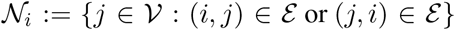 denotes the set of all in- and out-neighbors of node *i*. We denote the set of interior nodes, having no out-of-group neighbors by

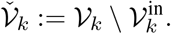

Lastly, we respectively denote the union of all boundary nodes in a partition by

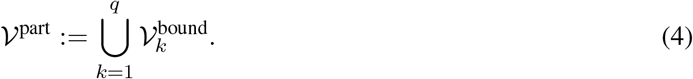

By intersecting LEQ sets on a node set *𝒱* ⊆ *𝒱*_0_, we form a set of *regional equilibrium* (REQ) states.

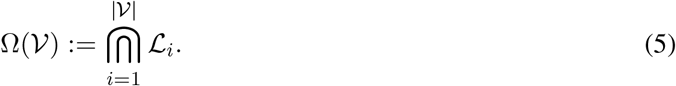

Let *r* be a free parameter that defines the maximum size of a node set that we will not partition further, but rather compute the full equilibrium set using (5). The set *Ƶ*(*𝒱*) represents the full state space of the nodes in *𝒱* and the empty set everywhere else:

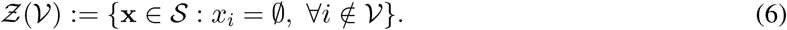

We are now ready to introduce the core of the proposed method, which is the following recursive definition of a compact representation of the REQ sets Ω(*𝒱*):

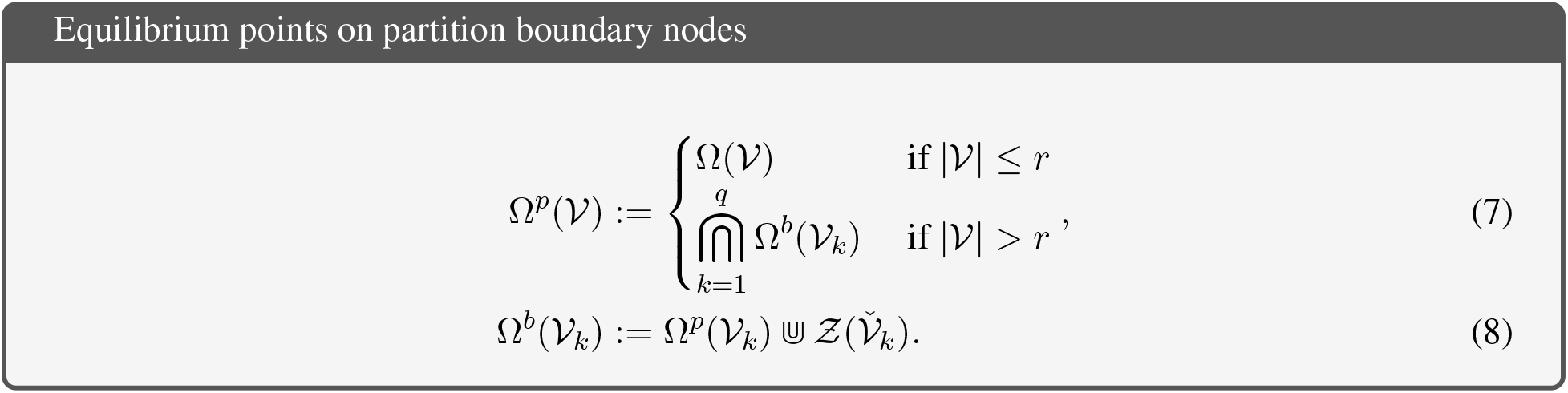

The set Ω^*p*^(*𝒱*) represents the set of all possible equilibrium values for nodes that lie on the boundary of the partition of 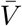. At the base level where |*𝒱*| ≤ *r*, the problem is small enough to solve by intersecting all contained LEQ sets as in (5), and Ω^*p*^(*𝒱*) is equal to the full regional equilibrium set. Otherwise, we obtain Ω^*p*^(*𝒱*) by intersecting the sets Ω^*b*^(*𝒱*) for each partitioned subgroup. Ω^*b*^(*𝒱*_*k*_) is the set of all possible regional equilibrium values at the nodes that lie on the boundary of *𝒱*_*k*_ with some other partitioned subnetwork *V*_*j*_, *j* ≠ *k*. This set is obtained by taking the union of the set Ω^*p*^(*𝒱*_*k*_) (defined recursively) with 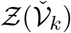. Taking the union of *Ƶ*(*𝒱*) with some sparsely represented set has the effect of decreasing the amount of information needed to represent the resulting set, since it will no longer contain any information about the nodes in *𝒱*. Notice that a graph partition 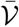 is needed to compute the lower expression in (7).

It is useful to store the results of this computation in the form of a tree graph 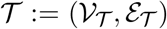, in which each node 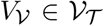 contains the partition 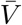, the set **Ω**^*p*^(*𝒱*) of possible equilibrium states on the partition boundary nodes, and the sets **Ω**^*b*^(*𝒱*_*k*_) and corresponding ∆ matrices for each partitioned subnetwork:

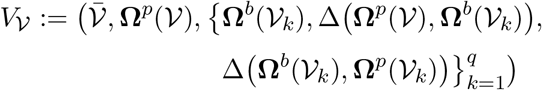

for each partition 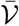 performed in the course of the recursion. The edges 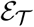 in the tree simply connect each tree node *V*_*𝒱*_ to the nodes corresponding to the partition of 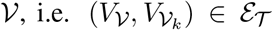 for each *k* ∈ {1, …, *q*}. As a result, we obtain a compact representation of the global equilibrium set. Specifically, Ω^*p*^(*𝒱*_0_) is equivalent to the union of the global equilibrium set with all state configurations of nodes interior to the top level partition, as formalized in the following theorem (proof in Appendix).

#### Theorem 2.

*For any partition* 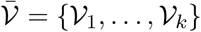 *of an arbitrary node set 𝒱 such that |𝒱*| > *r*,

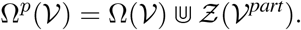

When paired with the information contained in 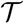, there is sufficient information in Ω^*p*^(*𝒱*) to efficiently compute the number of equilibria, the full equilibrium set, or only those equilibria that are optimal with respect to some cost function.

To provide further intuition into the approach, we illustrate a small example in Figure 6. The network shown in the center is partioned into groups indicated by the colored node outlines. Best-response dynamics with an anticoordinating payoff matrix of 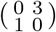 was used for all edges in the network (see **SI: Best response and linear threshold dynamics**). Two of the eight LEQ sets (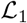 and 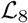) are shown towards the top, with the elements of the set in a stacked arrangement. The two PREQ sets (**Ω**^*p*^(*𝒱*_1_) and **Ω**^*p*^(*𝒱*_2_)) corresponding to the intersections of the LEQ sets within their respective partitioned subgroups are shown at the far left and right. Compact representations of these sets on their boundary nodes are shown in the BREQ sets (**Ω**^*b*^(*𝒱*_1_) and **Ω**^*b*^(*𝒱*_2_)) just underneath. Finally, the top level PREQ set **Ω**^*p*^(*𝒱*) resulting from the intersection of the two BREQ sets and the expanded global equilibrium set Ω(*𝒱*_0_) appear at the bottom.

**Figure 6:**
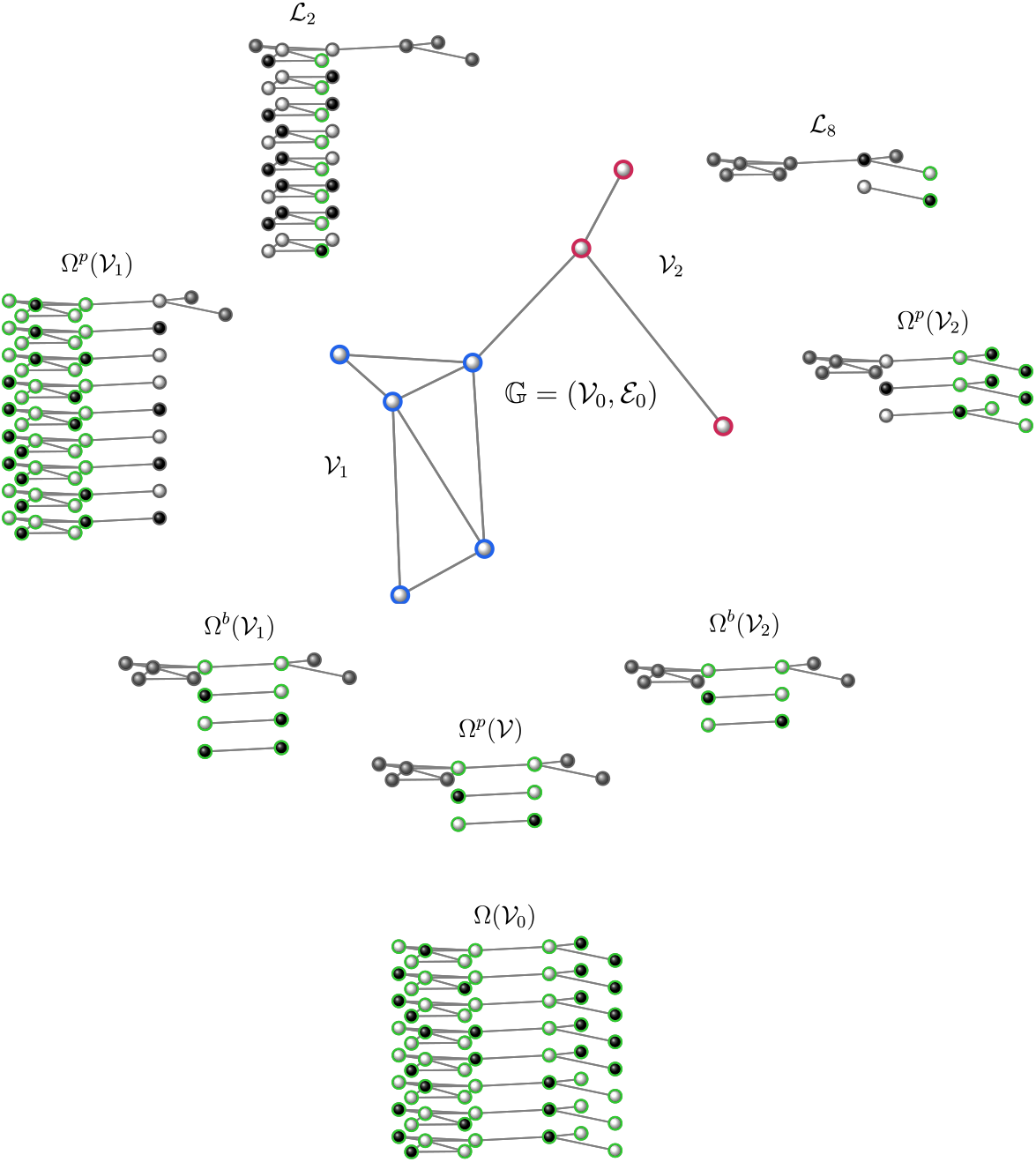
Example showing a network in which nodes can take one of two states marked by black or white (gray indicates nodes that can take any value), partitioned into two subnetworks *𝒱*_1_ (blue) and *𝒱*_2_ (red). We show two of the eight LEQ sets 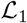 and 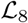. Below them are the sets Ω^*p*^(*𝒱*_1_) and Ω^*p*^(*𝒱*_2_), generated form the intersection of their constituent LEQ sets. Further below are the compactly represented sets Ω^*b*^(*𝒱*_1_) and Ω^*b*^(*𝒱*_2_) on the two partition boundary nodes. Finally, we show the top-level set Ω^*p*^(*𝒱*), and the global equilibrium set Ω(*𝒱*_0_) constructed from the information in 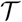.

### Existence and number of equilibria

The number of equilibria can be computed using the same recursion structure as above and can occur either simultaneous to searching the equilibrium space or in post-processing. To do this, at the lowest level of partitioning (the top case of (10)), we simply count in (9) how many regional equilibrium states are included in each element of the PREQ set. These quantities are then passed to the higher level in the partition tree along with the requisite bookkeeping steps, ultimately resulting in the total number of equilibria at the top level, as specified in the following theorem (proof in the **Appendix**).

#### Theorem 3.

*Given the tree* 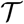 *resulting from the computation* (7)-(6) *on system* (1), *the total number of equilibria* |**Ω**(*𝒱*_0_)| *is given by*

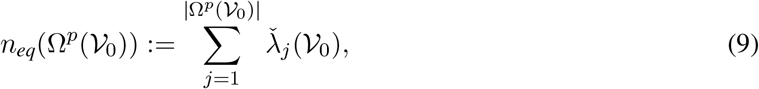

where

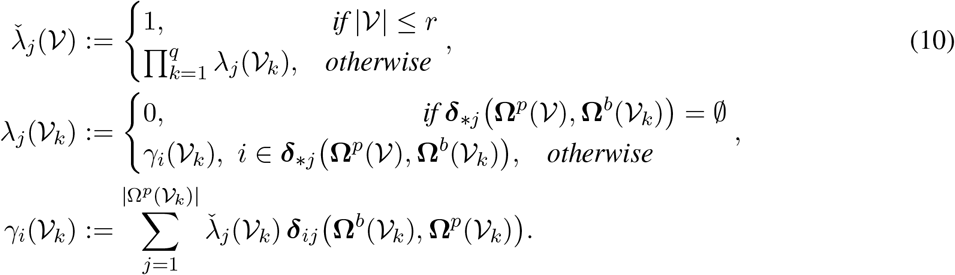

### Optimization on the global equilibrium set

Within the set of all fixed points, it is often useful to find the equilibria that optimize a given cost or energy function. An advantage of this approach is that the optimal equilibria can be quickly extracted from the results of the computation without generating the full equilibrium set, which can be very large in some cases.

Optimization using this approach works for the class of global cost functions arising from the linear superposition of possibly nonlinear local cost functions. Let 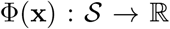 denote a cost function on the state space 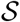, which is the summation of local node and edge costs:

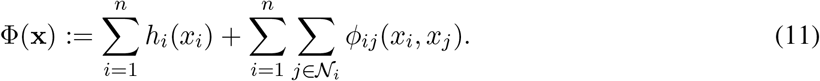

Suppose that we seek the set of equilibria that minimize this function:

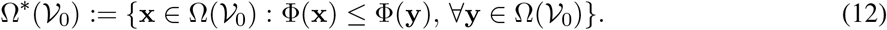

It is also useful to define the cost of a particular local or regional equilibrium set, which we denote by ***ω***(*𝒱*) *∈* Ω(*𝒱*), where Ω(*𝒱*):= *{**ω***_1_(*𝒱*), …, ***ω***_*α*_(*𝒱*)} and *α*:= |Ω(*𝒱*)|. Recalling that **x**[*i*] denotes the fixed state of node *i* in the sparse representation set 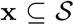, we note that ***ω***(*𝒱*)[*i*] has the same interpretation for the equilibrium set ***ω***(*𝒱*). We can now express the cost of ***ω***(*𝒱*) as follows:

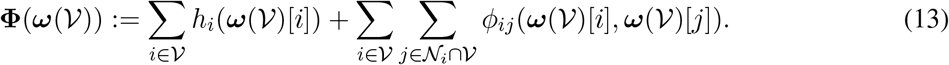

At the lowest level of the partition tree, the cost of each element ***ω***(*𝒱*) is computed according to (13). Notice that while the costs associated with all nodes of *𝒱* are included, only the costs of edges connecting pairs of nodes in *𝒱* are included, while costs related to edges containing nodes not in *𝒱* are excluded. These inter-region costs are and defined as follows:

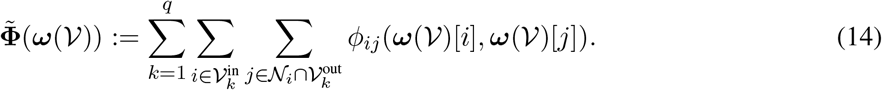

Although the cost of a base-level equilibrium set ***ω*** is unique and well-defined, the cost of a PREQ element ***ω***^*p*^(*𝒱*) or BREQ element ***ω***^*b*^(*𝒱*) may take multiple values, since they encapsulate the set of fixed points for all configurations of the interior nodes. Hence, we define the costs of these elements as ordered sets:

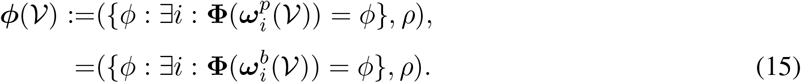

Note that the set of feasible cost values in the sets **Ω**^*p*^(*𝒱*) and **Ω**^*b*^(*𝒱*) are identical, since they include the same underlying REQ set Ω(*𝒱*). Next, we define a matrix that maps the elements of a PREQ set to the number of ways each unique cost value is attainable in the respective node groups, in which we have labeled the rows and columns for ease of interpretation:

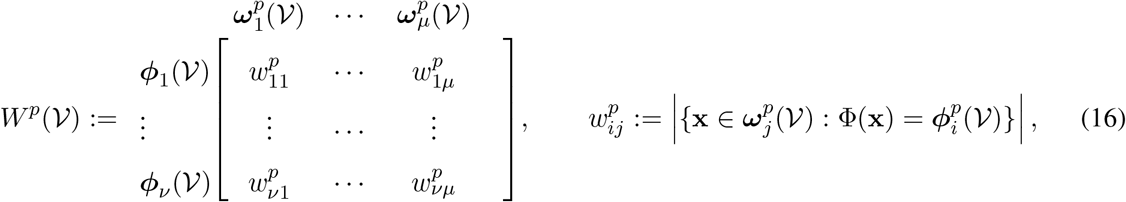

where *µ*:= |**Ω**^*p*^(*𝒱*)|, *ν*:= *|**φ***^*p*^(*𝒱*)|, and the matrix *W*^*b*^(*𝒱*) is defined similarly.

Clearly, computing the quantities (15)-(16) explicitly would require generating the entire set Ω(*𝒱*). Fortunately, there is a more efficient way to compute these using the partition tree structure 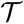 and the cost values attained by lower level REQ sets. To help with this task, it will be convenient to define two sets of matrices. First, the matrix 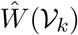 maps the unique costs achievable by elements of the lower level BREQ sets **Ω**^*b*^(*𝒱*_*k*_) to the current level PREQ elements ***ω***^*p*^(*𝒱*) with which the costs are compatible:

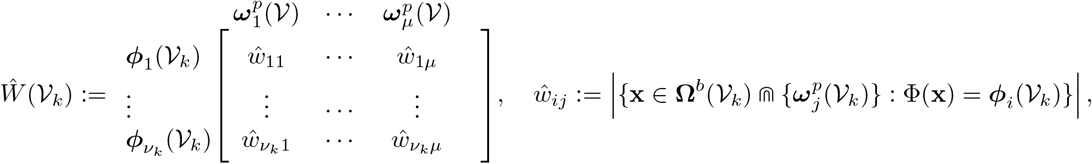

where *ν*_*k*_:= *|**φ***(*𝒱*_*k*_)|. Second, the matrix 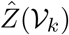 contains the costs associated with each entry in *Ŵ*(*𝒱*_*k*_), or ∞ if the entry corresponds to an infeasible pairing:

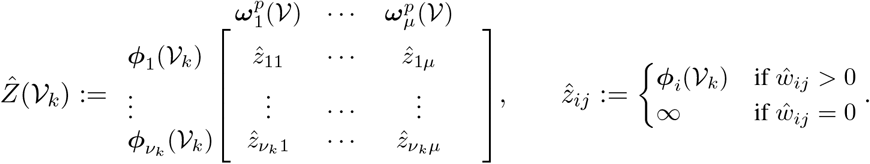

Next, we construct expanded matrices 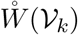 and 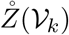 for each subgroup *k*. Let 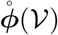 denote the sums of all configurations of the unique subgroup costs ***ϕ***(*𝒱*_*k*_), of which the total number is 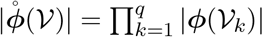. Then the expanded matrix 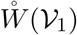 is given by:

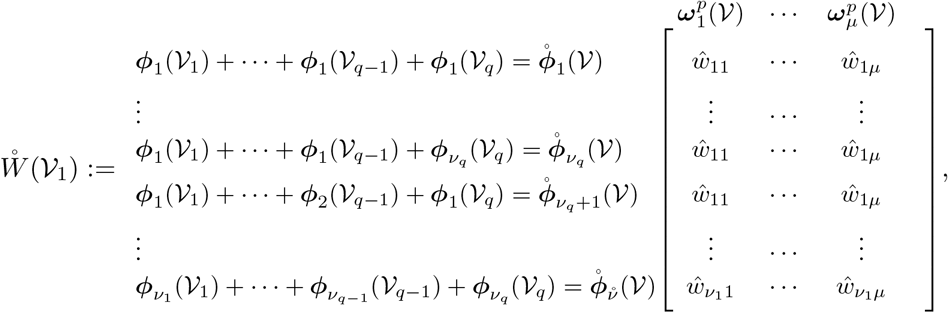

where *ŵ*_*ij*_ is equal to the *ij*^*th*^ entry of *Ŵ* (*𝒱*_1_). The successive matrices 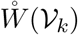 are defined similarly, with *ŵ*_*ij*_ appearing in the same row where ***ϕ***_*i*_(*𝒱*_*k*_) appears in the summations yielding the entries of 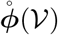. Likewise, the expanded matrix 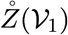 is given by:

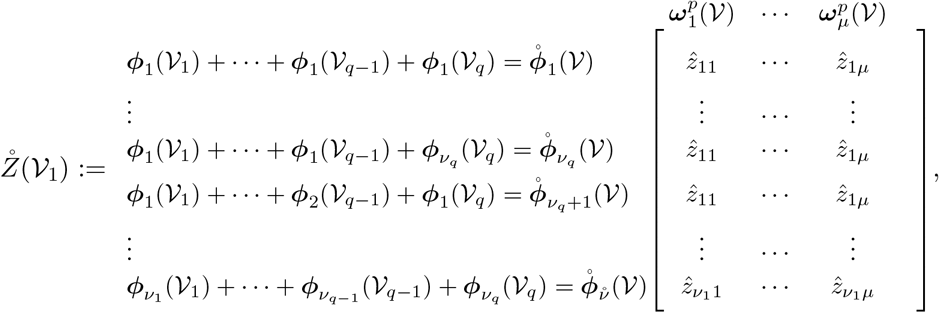

where 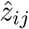 is equal to the *ij*^*th*^ entry of 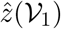. Next, let 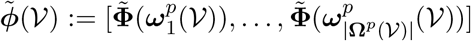 denote a row vector of inter-group costs. The combined expanded cost matrix 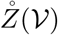 and the number of instantiations of each cost 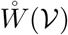 can be expressed as follows:

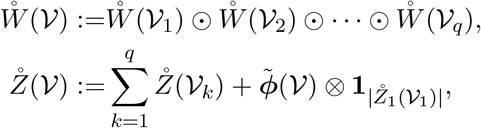

where ⊙ denotes element-wise multiplication and ⊗ denotes the Kronecker product. Finally, the compact matrix *W*^*p*^(*𝒱*) is constructed by extracting the unique cost values in 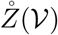 and counting the total number of ways each cost is attained in 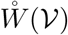. We again refer the reader to the Cost algorithm in **SI** for details on this procedure. The optimal equilibrium set Ω^*\**^(*𝒱*_0_) can then be obtained by following a similar procedure, as described in detail in the Expand algorithm (see **SI**). Note that by using the null cost function Φ(**x**) = 0, 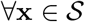, the set Ω^*\**^(*𝒱*_0_) is equivalent to the global equilibrium set Ω(*𝒱*_0_) of (1).

## Appendices

### Processing of data from Human Connectome Project

The data used consisted of resting state fMRI scans collected from 783 subjects during the Human Connectome Project (HCP). These subjects were part of the s900 release which contained a total of 818 subjects. However, 35 of these subjects which had been marked as having instabilities in the head coil were removed [45, 46]. Each subject underwent a total of four 15-minute scans divided among two separate scanning sessions. The two scans per session corresponded to a left-right and a right-left acquisition. Data was acquired at 3T with a temporal resolution (TR) of 720ms. Detailed information on HCP scan protocols are contained in [47].

The publicly available s900 data release includes the following processing procedure. Following the standard minimal preprocessing pipeline [47], the motion correction strategies recommended by Power and colleagues were applied [48], with the exception of Frame Censoring (removing time points), which the authors acknowledge has proven controversial. The remaining motion correction procedure consists of the FSL ICA-FIX correction [49] and regressing out the twelve HCP motion parameters and global signal (cerebrospinal fluid, white matter, and grey matter). To remove signal drift, we applied a .009 Hz high-pass filter and detrended to remove respiratory artifact. Data were parcellated into the Gordon atlas [50] and functional connectivities were computed using the Pearson correlation between parcels for the combined data across scans. More details on the task specializations associated with the various resting-state networks can be found in [51].

To suppress weak correlations and to facilitate comparison between subjects that are independent of overall connectivity variations, we thresholded correlation coefficients to preserve 5% of all pairwise connections, following similar methods as in [52]. Fixed points were computed for best-response dynamics in which the payoff matrices between each pair of regions *i* and *j* is given by *J*_*ij*_*I*_2_, where *J*_*ij*_ denotes the Pearson correlation coefficient between respective regions and *I*_2_ is the 2 *×* 2 identity matrix. These dynamics are guaranteed to converge to a local minimum of the Ising Hamiltonian:

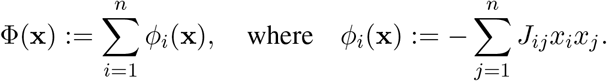

### Analysis of a coarse-grained lattice model for protein folding

For this case study, the recursive partitioning algorithm was augmented with two stages of trimming, one to eliminate infeasible entries in the LEQ sets, and a second to eliminate known infeasible entries in the intermediate REQ sets. These steps are possible due to the regularity of the lattice structure and the uniformity of the cost function across nodes, and are generally good practice when applicable because of the resulting computational savings.

### Proof of Theorem 1

*Proof.* The overall computation involves three types of operations: local equilibrium checks, sparse set intersections, and computing the ∆ mappings.

#### Computing the local equilibria

Finding the local equilibrium states at each node involves evaluating the local update rule for a maximum of 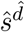 total state configurations. Multiplying by the number of nodes *n* yields a computational complexity of 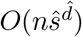 for this step.

#### Sparse set intersections

The sparse set intersections occur at two levels. At the base level, the intersection of a pair of local equilibrium sets involves checking a maximum of 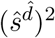 overlapping state configurations. The result is then checked against a third local equilibrium set, and the process repeats until the maximum group size *r* is reached, yielding a total for each base-level partition group of

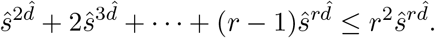

Assuming maximum group sizes, this computation is performed approximately 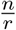 times, yielding an upper bound of 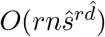 for the local intersections.

Intersections at all higher levels depend on the number of boundary nodes between neighboring partition groups, of which there are a maximum of 2*k*. However, the states of all boundary nodes must be represented for each partition group. Since there up to *q* partition groups, all of which may be adjacent to each other, the maximum dimension in each group is 2*kq*. Hence, each intersection involves checking up to (*ŝ*^2*kq*^)^2^ overlapping state configurations, each of which may contain up to 2*kq* nodes, resulting in a complexity of *O*(2*kqŝ*^4*kq*^) for each partition that is performed. Since the total number of partitions is bounded above by *n*, we have a total complexity of *O*(*kqnŝ*^4*kq*^) for the higher level sparse set intersections. The computational complexity of all sparse set intersections is then no greater than 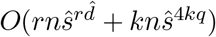.

#### Computing the ∆ matrices

The ∆ matrices defined in (2) in the main article map regional equilibrium states between neighboring levels in the partition hierarchy. Since the maximum number of configurations on each level is *O*(*ŝ*^2*kq*^), the number of entries in each ∆ matrix is at most *O*(*ŝ*^4*kq*^). Since each entry *δ*_*ij*_ requires at most 2*kq* computations (see (3) in the article), the computational complexity for constructing each ∆ matrix is bounded by *O*(*kqŝ*^4*kq*^). Finally, since both the number of nodes and edges in the tree 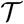 are less than *n*, the number of ∆ matrices to compute is at most 2*n*, resulting in a complexity of *O*(*kqnŝ*^4*kq*^).

The overall computational complexity of computing 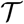 is therefore bounded by 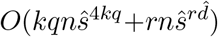.□

### Proof of Theorem 2

*Proof.* Since *|𝒱*| > *r*, according to (7) and (8),

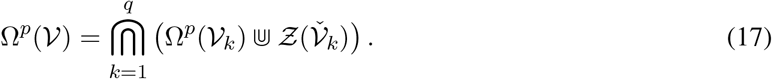

Pulling out the first term in this expression yields

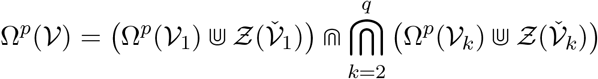

which can be expanded to

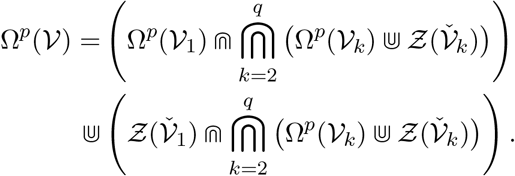

Due to (6) and the fact that the partition is disjoint, we know that 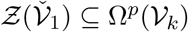 and that 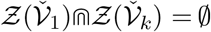 for all *k* ∈ {1, …, *q*}. It follows that

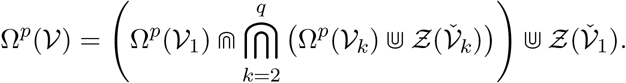

Noticing that the middle term is in the same form as (17), we can repeatedly apply the previous two steps to obtain

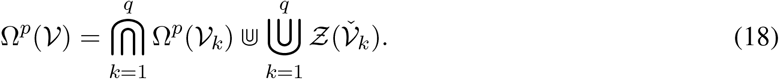

Next, suppose that |*𝒱*_*k*_| > *r* for each *k* and let 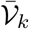 denote a partition of each *𝒱*_*k*_. We can then perform the exact same expansion of each Ω^*p*^(*𝒱*_*k*_) from (17) to (18), resulting in

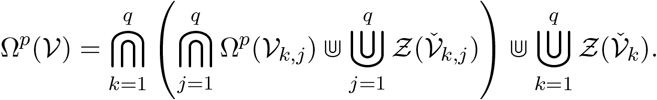

Similarly as above, since 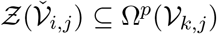 and 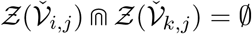 for all *i, j, k* ∈ {1, …, *q*}, *i* ≠ *k*, we have

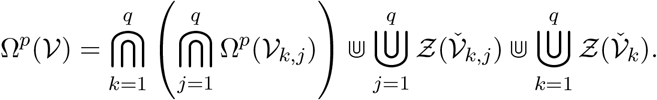

Since 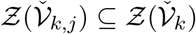 for each *j* ∈ {1, …, *q*}, we can simplify as follows:

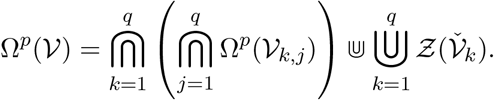

By repeatedly performing this expansion according to (8), the term on the left will eventually become the union of base-level regional equilibrium sets **Ω**(*𝒱*_*k,j*,…_), which is equal to **Ω**(*𝒱*). Furthermore, since 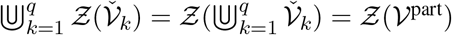, due to (4), we obtain

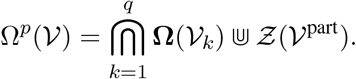

Finally, since

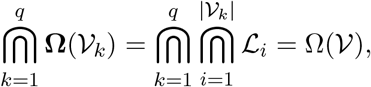

we have the desired result 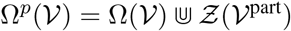 and the proof is completed. □

### Proof of Theorem 3

*Proof.* Suppose *|𝭁*| ≤ *r*. Then

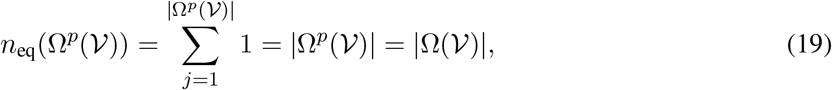

where we used the fact that |Ω^*p*^(*𝒱*)| = |Ω(*𝒱*)| from (7). Next, suppose *|𝒱*| > *r*. Then we have

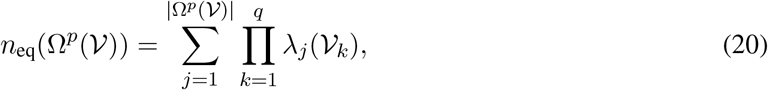

where |Ω^*p*^(*𝒱*)| is the number of unique configurations of partition boundary nodes in *𝒱* that appear in the global equilibrium set, as proved in Theorem 2. Given one of these configurations, the regional equilibria in each group are independent from each other. Therefore, it suffices to show that *λ*_*j*_(*𝒱*_*k*_) is equal to the number of REQ states associated with the set *𝒱*_*k*_ that are contained in 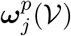, i.e. 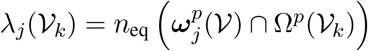.

We can check for compatibility between each entry of Ω^*p*^(*𝒱*) and the constituent BREQ states Ω^*b*^(*𝒱*_*k*_) using ***δ***_\**j*_(Ω^*p*^(*𝒱*), Ω^*b*^(*𝒱*_*k*_)). If there exists *j* such that this set is empty for some *k*, then *λ*_*j*_(*𝒱*_*k*_) = 0 and there are no equilibria. Otherwise, ***δ***_\**j*_(Ω^*p*^(*𝒱*), Ω^*b*^(*𝒱*_*k*_)) is a singleton since Ω^*b*^(*𝒱*_*k*_) is defined on a subset of the nodes defined in Ω^*p*^(*𝒱*). One can then check that the quantity *γ*_*i*_(*𝒱*_*k*_) is precisely equivalent to 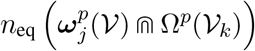, which is then equal to |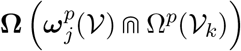 | if |*𝒱*_*k*_| ≤ *r*, and otherwise the number of compatible lower-level equilibria, and the proof proceeds by induction.□

#### Best response and linear threshold dynamics

A common type of network dynamics involves nodes that optimize local cost or *utility* functions. For our purposes, these dynamics are particularly useful in constructing systems that will converge to fixed points and local minima of a function. Let *u*_*i*_(*x*_*i*_, **x**_*−i*_) denote the utility to node *i* when the states of all nodes other than *i* are collected in the vector **x**_*−i*_. A simple best-response update is given by

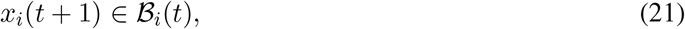

where the set of best responses is

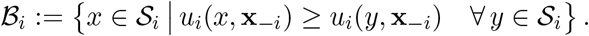

In particular, suppose that each node can take one of two possible states, i.e. 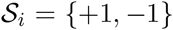 for all *i* ∈ *𝒱*, and the utility of each node can be expressed as the sum of outcomes of 2 *×* 2 matrix games defined on each outgoing edge from node *i*:

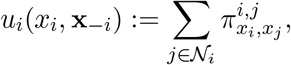

where the *payoff matrix π*^*i,j*^ defines the outcome of a game in which node *i* chooses the row and node *j* chooses a column:

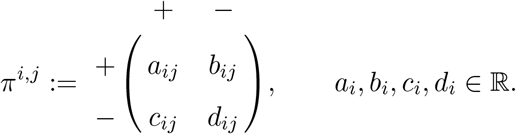

The utility can then be expressed as *u*_*i*_(*x*_*i*_, **x**_*−i*_) =

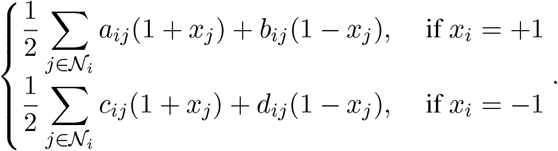

The best-response update (21) now becomes

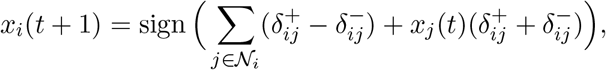

where 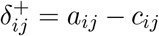 and 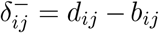. Further simplification reveals a type of linear threshold model, in which a weighted sum of neighbor states is compared to a local threshold value *τ*_*i*_:

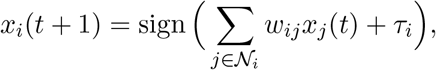

where 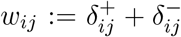 and 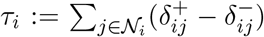. Threshold models are prominent in various research fields, from computational neuroscience [1] to sociology [53]. A remarkable property of these models in addition to their ability to describe wide-ranging physical phenomena is that, when nodes update asynchronously, the networks will converge to a fixed point if any of the following conditions hold:

1. all weights (payoff matrices) are symmetric, i.e. *w*_*ij*_ = *w*_*ji*_ (*π*^*ij*^ = *π*^*ji*^), for all *i, j* ∈ *𝒱* [1];
2. all neighbors are treated equally, i.e. *w*_*ij*_ = *w*_*ik*_ (*π*^*ij*^ = *π*^*ik*^) for each *i* ∈ *𝒱* and all *j*, 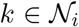, and all nodes are *coordinating*, i.e.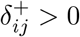 and 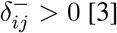 [3];
3. all neighbors are treated equally and all nodes are *anticoordinating*, i.e. 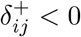 and 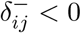 [3].

#### Graph partitioning and separability analysis

Although it is computationally complex to determine the partition separability of networks in general, there are several graph partitioning algorithms that try to minimize the number of edges between adjacent partition groups while producing groups of approximately equal size [17, 44]. To determine which types of networks are best suited to the proposed approach, we recursively applied the algorithm in [17] on three standard network topologies: geometric random networks (constructed as in **Computational complexity simulations**), small-world networks (constructed using the Watts-Strogatz method [54] with four nearest-neighbor connections and rewire probability 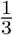), and scale-free networks (constructed using preferential attachment with a minimum degree of three [55]). Fig. 7 shows the ratio of the maximum number of edges crossing partition boundaries to the total number of edges in 100 random networks for each of these types. We observe that geometric random networks and small world networks tend to be separable along a smaller portion of edges than scale-free networks, and are therefore better suited to the methods proposed in this article.

**Figure 7:**
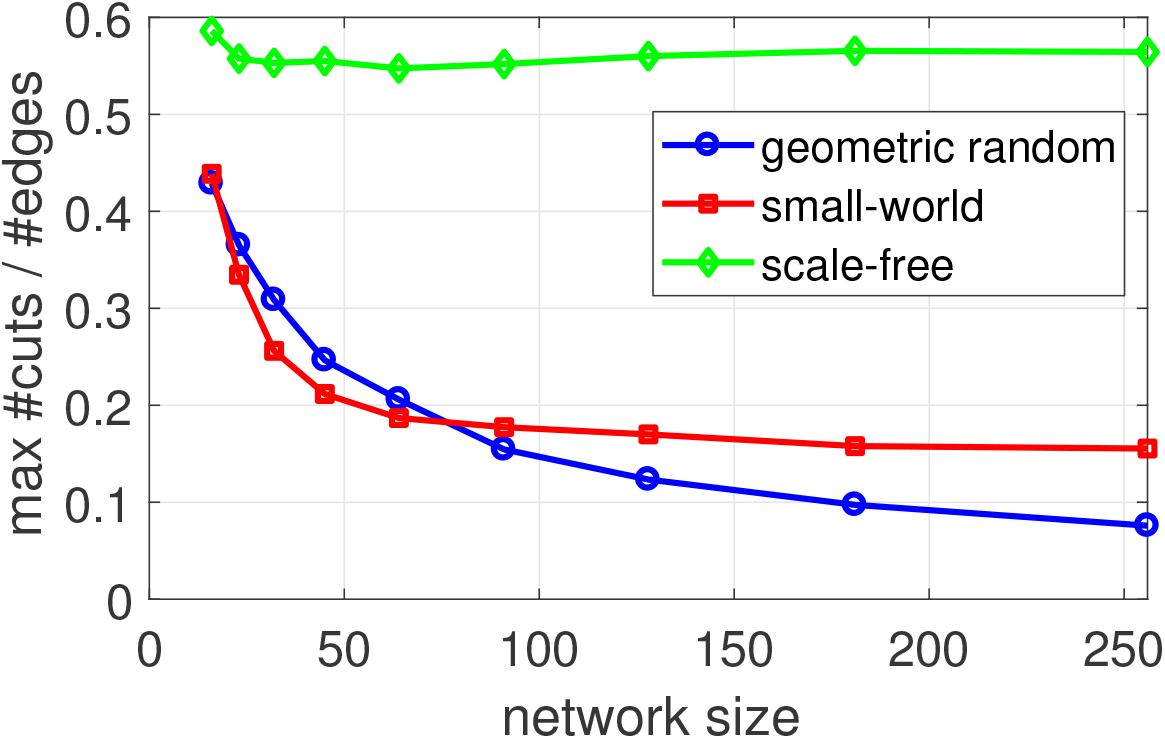
Maximum number of edges crossing partition boundary for 100 randomly generated networks of each type (geometric random, small-world, scale-free).

#### Computational complexity simulations

The computational complexity study shown in Fig. 1 in the main article consisted of three different network topologies (ring networks, tree networks, and geometric random networks), governed by the bestresponse threshold dynamics described above. Energy was computed using (11) with *ϕ*_*ij*_(*x*_*i*_, *x*_*j*_):= *x*_*i*_*x*_*j*_ and *h*_*i*_(*x*_*i*_):= 0 for each node *i* ∈ *𝒱*.

The ring networks were undirected and anticoordinating payoff matrices of 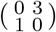 were used for all edges. Undirected tree networks used in Fig. 1 were constructed by starting from a root node and randomly adding between 0 and 4 branches with uniform probability (0.2), and repeating the process at each of the leaf nodes for up to a maximum number of generations, which was increased from 1 to 11. Coordinating payoff matrices of 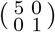 were used for all edges.

Geometric random networks were constructed with sizes *n*:= ⌈2^*κ*^⌉ for each *κ* ∈ {3, 3.5, 4, 4.5, 5, 5.5, 6, 6.5, 7, 7.5, 8, 8.5, 9} by fixing a mean degree 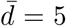 and then setting the connection radius for each size *n* to 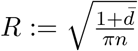. After uniformly randomly distributing *n* nodes in the unit square, edges were added between every pair of nodes within a distance *R* of each other. Any disconnected networks were discarded. Coordinating payoff matrices of 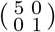 were used for all edges.

## Algorithm pseudocode

To augment the mathematical framework provided in the previous sections, we provide here a step by step implementation of the main algorithm in pseudocode. Due to their length, we provide the algorithms for computing the cost and expanding the optimal or full equilibrium set in the **Appendix**.

### Algorithm

Main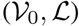

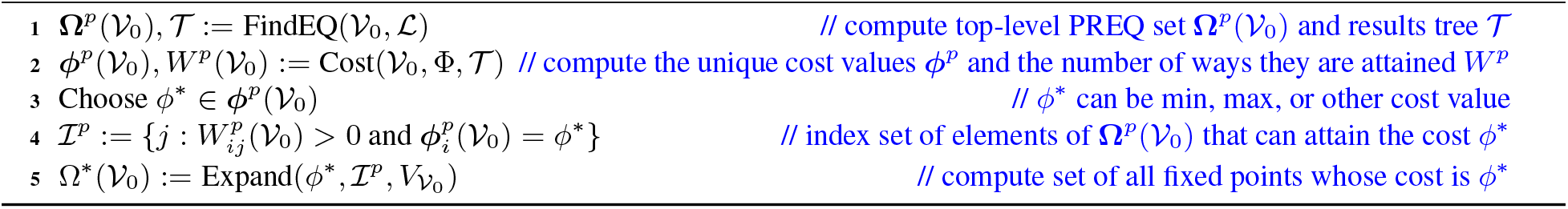

### Algorithm

FindEQ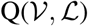

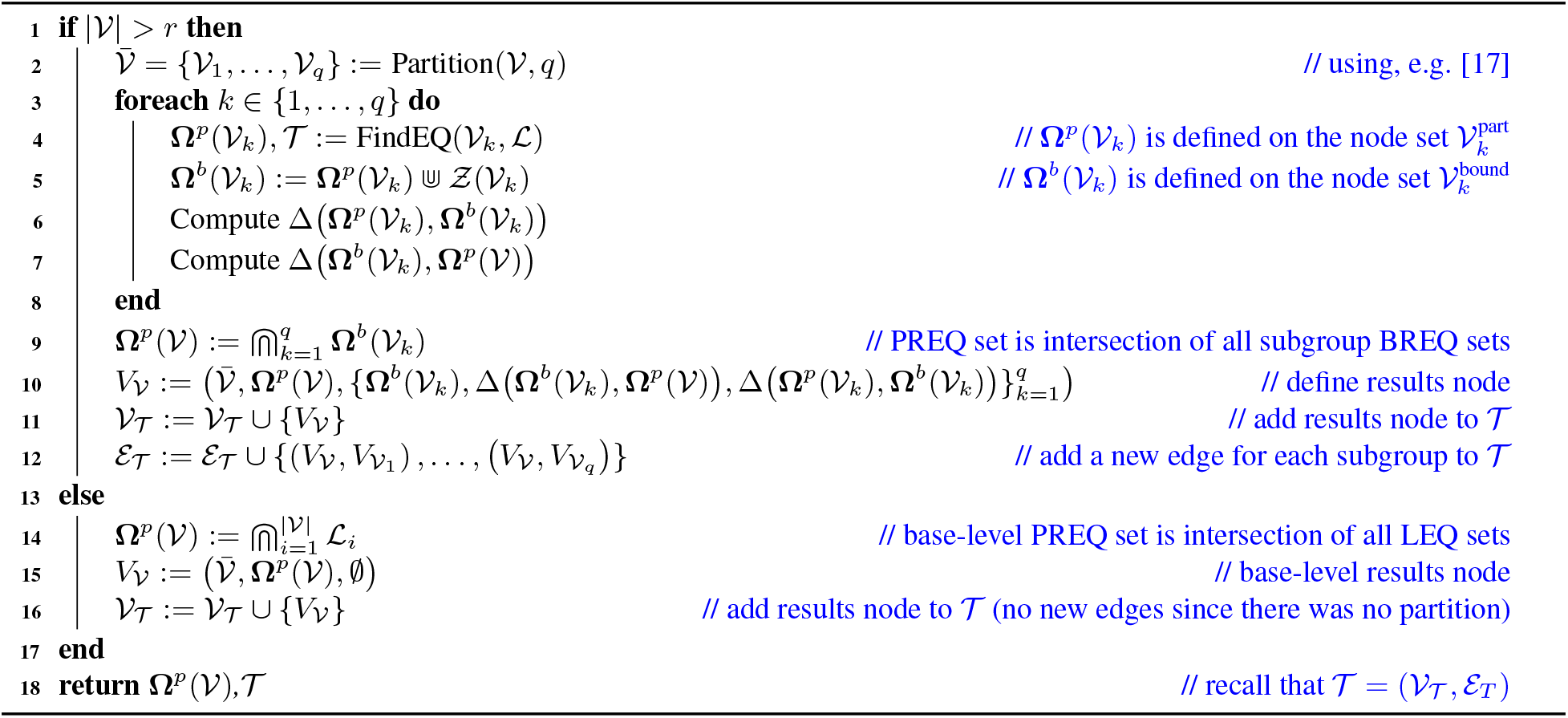

### Cost and Expansion Algorithms

Here we provide pseudocode implementations of the algorithms for computing the costs and expanding the optimal or full equilibrium sets.

#### Algorithm

Cost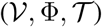

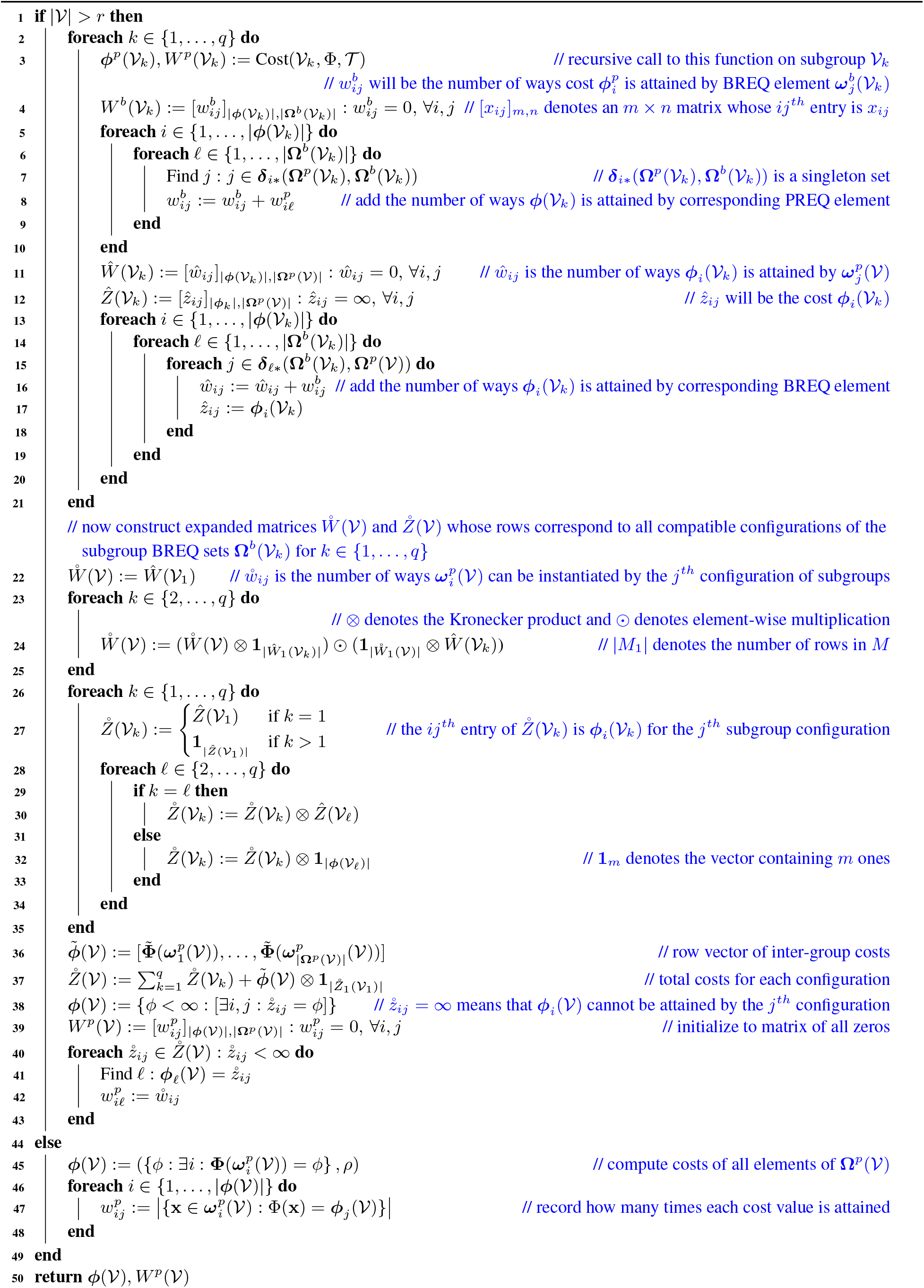

#### Algorithm

Expand 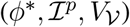

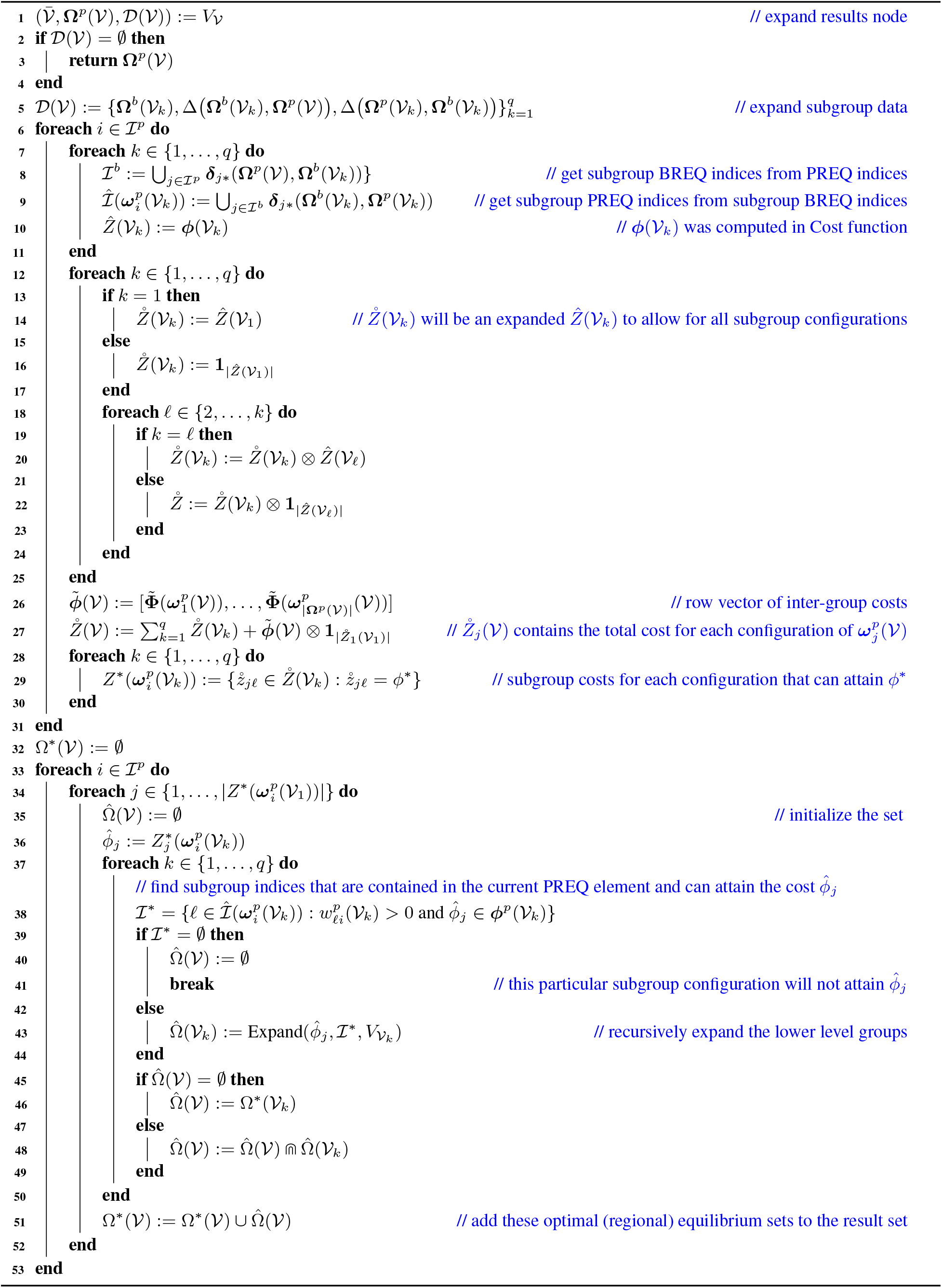

